# Impaired 26S proteasome causes learning and memory deficiency and induces neuroinflammation mediated by NF-κB in mice

**DOI:** 10.1101/2024.02.09.579699

**Authors:** Christa C. Huber, Eduardo Callegari, Maria Paez, Xiaoping Li, Abena Dwamena, Hongmin Wang

## Abstract

A reduction in proteasome activity, loss of synapses, and increased neuroinflammation in the brain are hallmarks of aging and many neurodegenerative disorders, including Alzheimer’s disease (AD); however, whether proteasome dysfunction is causative to neuroinflammation remains less understood. In this study, we investigated the impact of 26S proteasome deficiency on neuroinflammation in the *Psmc1* knockout (KO) mice deficient in a 19S proteasome subunit limited to the forebrain region. Our results revealed that impaired 26S proteasome causes reduced learning and memory capability and overt neuroinflammation in the *Psmc1* KO brain at eight weeks of age. Pronounced neuroinflammation was confirmed by increased levels of several key immune response-related proteins, including Stat1, Trem2, and NF-κB, and by activation of astrocytes and microglia in the KO brain. To validate NF-κB mediating neuroinflammation, we administered a selective NF-κB inhibitor to the KO animals, and following the treatment, the KO mice exhibited improved learning and memory behaviors and reduced neuroinflammation compared to the control animals. These data indicate that impaired 26S proteasome causes AD-like cognitive deficiency and induces neuroinflammation mediated largely by NF-κB. These results may aid the development of effective therapeutics for AD and other neurodegenerative disorders where impaired proteasome activity is consistently coupled with neuroinflammation.

## Introduction

Misfolded, aberrant, or unneeded intracellular proteins are degraded by the ubiquitin-proteasome system (UPS), a major protein degradation system of the cell that functions to maintain intracellular proteostasis. Many studies have shown that proteasome activity decreases in the patient’s brain of Alzheimer’s disease (AD) and many other neurodegenerative diseases (Keller *et al*, 2000; Li & Guo, 2009; Liu *et al*, 2014; Medina *et al*, 2011; Wang & Wang, 2020). AD, an age-related disease, causes neurodegeneration in the brain, which can be attributed to robust neuroinflammation and impaired proteasome function (Guzman-Martinez *et al*, 2019).

Neuroinflammation is an inflammatory response in the nervous system that is considered to be neuroprotective initially in the brain by eliminating diverse pathogens following microbial infections (Tran *et al*, 2022), boosting tissue repair, and removing debris; however, increasing evidence has suggested that sustained neuroinflammation is detrimental to neuronal function and survival (Kwon & Koh, 2020). Many endogenous stimuli, such as genetic mutation and protein accumulation and aggregation, can persist in activating microglia and astrocytes to transition into proinflammatory phenotypes and causing neurodegeneration (Wei & Li, 2022; Zhao *et al*, 2017). Nevertheless, how neuroinflammation occurs in AD and other neurodegenerative diseases where proteasome function is impaired remains less understood.

For a protein to be degraded by the UPS, it needs to be tagged by a polyubiquitin chain through a series of repeated processes known as polyubiquitination. The substrate protein can then be recognized and degraded by the proteasome. The proteasome is a large ATP-dependent complex containing the 20S proteolytic core particle associated with either one or two 19S regulatory particles to make up the 26S or 30S proteasome, respectively. The 20S core particle is arranged in an α7-β7-β7-α7 fashion (Fig. 1A). The α-rings allow unfolded proteins to enter where the β-rings are located. The proteolytic activities of the 20S proteasome core are determined by the three β subunits of the proteasome: the β1 subunit offering caspase-like activity, the β2 subunit having trypsin-like activity, and the β5 subunit possessing chymotrypsin-like activity.

**Fig. 1.**
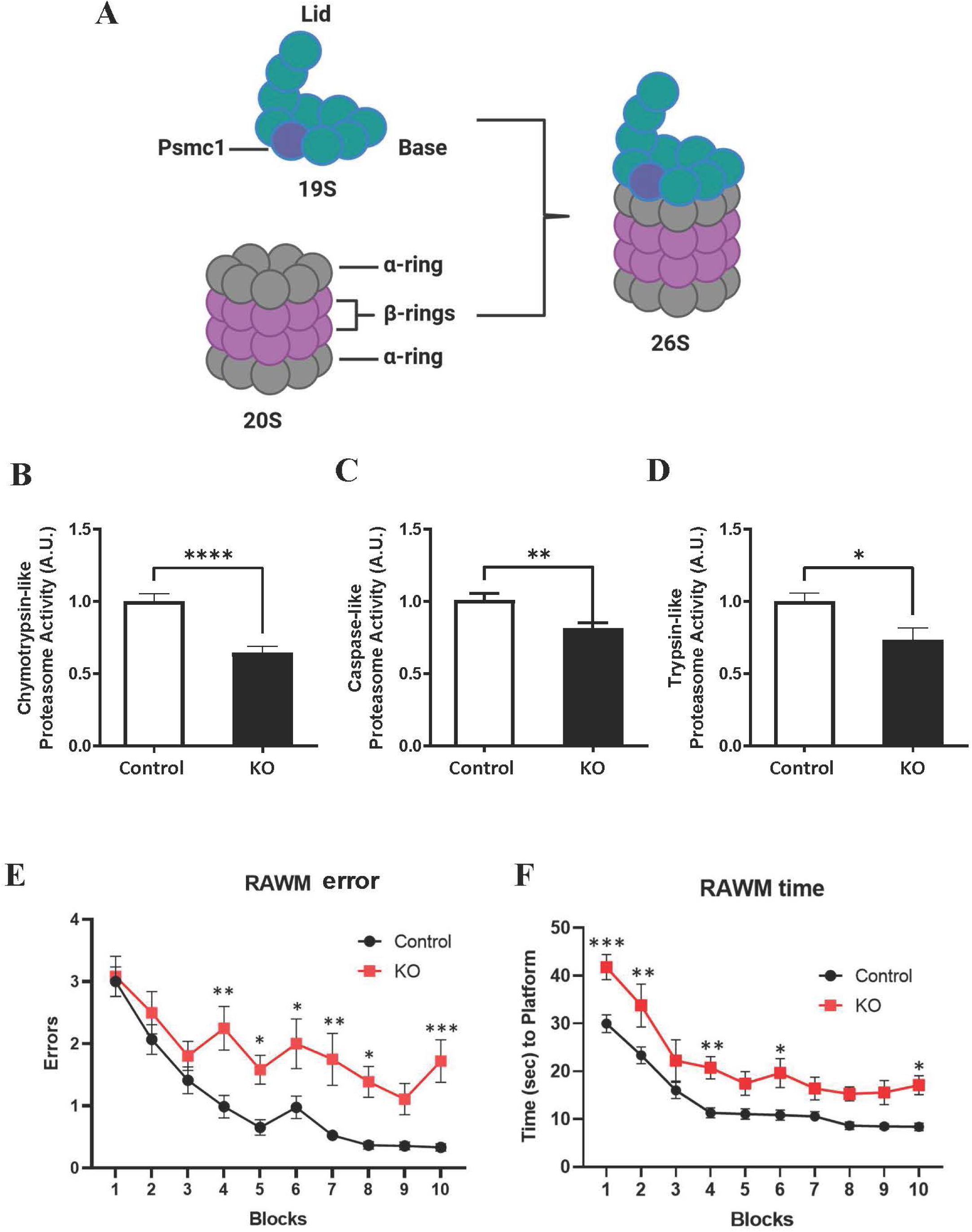
Genetically disrupting 26S proteasome reduces proteasome activity and impairs animal learning and memory. A diagram illustrating the structures of both the 20S and 19S proteasomes (**A**). The subunit, Psmc1, of 19S, whose gene was disrupted in this study, was highlighted and pointed out. Chymotrypsin-(**B**), caspase-(**C**), and trypsin-like (**D**) proteasome activities were reduced in the forebrain of the KO mice (n = 6, 3 males and 3 females for each genotype). The RAWM was used to assess animals’ spatial learning and memory by counting the number of incorrect arm entries (errors) (**E**) and the time it takes each mouse to find the platform (**F**). Data are presented as mean ± SEM; n = 12 - 29 per genotype. * p < 0.05, ** p < 0.01, *** p < 0.001, **** p < 0.0001 (RAWM data analyzed by repeated measures, two-way ANOVA, and Sidak’s multiple comparisons test. All other data was analyzed by a student’s t-test).

Structurally, the 19S regulatory particle is divided into two parts, the lid, and base (**Fig. 1A**). The base has six AAA+ ATPases and four non-ATPases. While the non-ATPases function in ubiquitin recognition and binding, the AAA+ ATPases assist in unfolding the misfolded protein and in opening the α-subunits to translocate the unfolded protein into the 20S core. To assemble into a functional 19S regulatory particle, Rpn1, Rpn2, and Rpt1-6 are needed. It is noted that Rpt2, also known as *Psmc1*, plays a key role in opening the entry pore gated by the α-subunits of the 20S proteasome core, allowing a substrate protein to enter the catalytic chamber for turnover (Budenholzer *et al*, 2017; Ding & Shen, 2008; Fernandez-Cruz & Reynaud, 2021). In the brain, the proteasome plays a crucial role in the removal of misfolded or unwanted proteins to maintain normal cognitive function, synaptic plasticity, signal transduction, and immune responses (Collins & Goldberg, 2017; Ding & Shen, 2008; Gupta *et al*, 2018).

Impaired proteasome and neuroinflammation are two major characteristics of aging, contributing to a variety of age-related neurodegenerative diseases, including AD. Aging is closely associated with protein misfolding, proteasomal dysfunction, and protein aggregation. On the other hand, accumulating data have shown that reduced proteasome activity is an early event of brain aging and a driver of neurodegeneration (Kelmer Sacramento *et al*, 2020). Given the fact that reduced proteasome activity and increased neuroinflammation are two common pathological features, or hallmarks, of aging, AD, and many other neurodegenerative disorders, we hypothesize that 26S proteasome deficiency is a causative mechanism to induce neuroinflammation in the brain. To test this hypothesis, we genetically ablated *Psmc1*, one 19S proteasomal subunit of the 26S proteasome, in neurons of the forebrain region. This genetic ablation impairs the assembly and activity of the 26S proteasome in this brain region. Utilizing this model, we discovered that 26S proteasome deficiency drives neuroinflammation and AD-like cognitive behaviors mediated by the NF-κB signaling pathway, while inhibition of the NF-κB pathway attenuates neuroinflammation and cognitive impairment.

This work is based on Chapter 3 of the first author’s PhD dissertation (https://red.library.usd.edu/cgi/viewcontent.cgi?article=1128&context=diss-thesis).

## Materials and Methods

### Animals

All procedures used to handle mice were approved by the Institutional Animal Care and Use Committee at the University of South Dakota and were in accordance with the National Institute of Health Guide for the Care and Use of Laboratory Animals. Mice were housed in four animals/cage on a 14/10-hour light/dark cycle. Mice were allowed to reach water and food ad libitum. Floxed *Psmc1* mice were previously described (Bedford *et al*, 2008), and CamKIIα-Cre (T29-Cre) transgenic mice were obtained from the Jackson Laboratory (stock #: 005359). The breeding strategy was according to the breeding recommendation of Jackson Laboratory to generate *Psmc1* KO (fl/fl)/T29-Cre (Tg/wt) animals in the forebrain. Both male and female mice were used for experiments at two months of age. The experiments were carried out with the experimenters blinded to the genotypes of the mice.

The *Psmc1* gene encodes for the Rpt2 subunit of the 19S regulatory particle. The CamkIIα-Cre mice target expression to neurons in the forebrain region of the brain (Tsien *et al*, 1996); therefore, ubiquitin-dependent degradation of proteins will be prevented in neurons in the forebrain region of the mice carrying both the Cre and homozygous floxed *Psmc1* gene (Bedford *et al*., 2008).

### Animal learning and memory test

The radial arm water maze (RAWM) was used to measure the learning and memory capability of mice as previously described (Adegoke *et al*, 2017; Alamed *et al*, 2006). Briefly, each mouse was gently placed into an arm and allowed to find the platform located in the goal arm. On day 1, mice were trained for 15 trials. The visible and hidden platforms were alternated until the 12^th^ trial, and the final 3 trials were performed using the hidden platform. On day 2, all 15 trials were performed with the hidden platform. Both the number of incorrect arm entries (errors) and the time spent to find the platform were recorded, using 60 seconds as a cut-off. The RAWM consists of 30 trials, with 3 consecutive trials being averaged to denote a block for a total of 10 blocks.

### Treating mice with NF-κB inhibitor

*Psmc1* KO mice were treated with pyrrolidine dithiocarbamate ammonium (PDTC), a selective NF-κB inhibitor (Chu *et al*, 2014), daily by intraperitoneal injections at a dose of 50 mg/kg for three weeks once mice reached five weeks of age. At eight weeks of age, mice were subjected to the RAWM test described above. On the day of behavioral tests, injections were performed early in the morning, and behavioral tests began at midday. Biweekly body weights of the mice were taken to monitor for any side effects of the drug.

### Proteasome activity assay

Proteasome activity assay was performed according to previously described methods (Liu *et al*, 2017). Briefly, the brain cortex was lysed in proteasome activity assay buffer (50 mM Tris-HCl, pH 7.5, 250 mM sucrose, 5 mM MgCl_2_, 0.5 mM EDTA, 2 mM ATP, and 1 mM dithiothreitol) by passing them ten times through a 27-gauge needle attached to a 1 ml syringe. After protein quantitation, 20 µg of total proteins from the lysates was used for detection. The fluorogenic substrate Suc-Leu-Leu-Val-Tyr-AMC (40 µM) was used to measure the chymotrypsin-like activity of the proteasome. Z-Leu-Leu-Glu-AMC (40 µM) was used to measure the caspase-like activity of the proteasome and the Boc-Leu-Arg-Arg-AMC (40 µM) was used to measure the trypsin-like activity of the proteasome. Fluorescence intensity was measured at 380 nm excitation and 460 nm emission using a plate reader (PerkinElmer, Waltham, MA, USA).

### Synaptosome isolation

Synaptosome isolation was performed according to previously described protocols (Hajos, 1975). Mouse brain cortex was homogenized on ice in 7 mL of 0.32 M sucrose containing protease inhibitor cocktail, 10 mM DDT, and deubiquitinase inhibitors (5 mM *N*-ethylmaleimide and 50 mM iodoacetamide) in a 2-mL dounce homogenizer for ten strokes. After centrifugation at 4°C for 10 minutes at 1,000 × *g*, the supernatant of the homogenates was transferred on top of an equal volume of 1.2 M sucrose in ultracentrifuge tubes and ultracentrifuged at 4°C for 35 minutes at 160,000 × *g* in an SW41 rotor in the Optima XPN-100 Beckman Coulter Ultracentrifuge. The synaptosome fraction from the cloudy interface between 0.32 M and 1.2 M sucrose was collected.

### Mass spectrometric analysis of synaptic proteins

The sample analysis was performed based on previously described methods (Huber *et al*, 2022; Paez & Callegari, 2022) (also see https://red.library.usd.edu/cgi/viewcontent.cgi?article=1128&context=diss-thesis). Briefly, the proteins were separated using 12% SDS-PAGE, and the gel was stained with colloidal Coomassie G250 (BioRad) and destained overnight. Each lane of the gel corresponding to each replicate/group was cut into gel slices. Each gel slice was destained, followed by reduction, alkylated, and subjected to in-gel digestion before being analyzed in an Easy nLC 1200 using a trapping and desalting online trap column (300 μm x 20 mm Acclaim PepMap C18 100Å (Thermo Scientific), and separated by an Easy-Spray PepMap RSLC 2 μm, 75 μm x 15 cm, nanoViper (Thermo Scientific) coupled to nanoESI orbitrap Exploris 240 HR/MA mass spectrometer. The chromatography conditions were as follows: 0–1 min, 1 % B isocratic; 2–60 min, 1–25%, 61-90 min 25-50%, and 92-101 min 50-100% B linear. Mobile Phase A (Water/Formic Acid, 99.9:0.1% v/v), and Phase B (Water/Acetonitrile/Formic Acid, 20/80/0.1% v/v). The solvent flow rate was 300 nL per minute. The orbitrap Exploris 240 instrument was operated under dependent mode (DDS), which is the top 12 modes to automatically switch between full scan MS and MS/MS acquisition. The ions were analyzed in positive MS ion mode (m/z 375 −1500) with 120,000 resolution (m/z 200) after accumulation with target ions to 1 × 106 value based on predictive AGC. The MS/MS ions selection was set at m/z 65-1800 to 1 × 105 counts. The spectrum was deconvoluted and analyzed using the Mascot Distiller v2.6 (www.matrixscience.com) and Proteome Discoverer v2.5 (Thermo Scientific). A mascot generic format list (MGF format) was generated to identify +1 or multiple charged precursor ions from the MS data file.

Mascot server v2.8.1 (www.matrix-science.com, UK) in MS/MS ion search mode (local licenses) was applied to conduct peptide matches (peptide masses and sequence tags) and protein searches against SwissProt Mus musculus 092022 (17562 sequences; 9,985,721 residues) database. The following parameters were set for the search: carbamidomethyl (C) on cysteine was fixed; variable modifications included asparagine and glutamine deamidation and methionine oxidation. Two missed cleavages were allowed; monoisotopic masses were counted; the precursor peptide and fragment mass tolerance were set at 15 ppm and 0.02 Da, respectively; the ion score or expected cut-off was set at 5. The MS/MS spectra were searched with MASCOT using a 95% confidence interval (% C.I.) threshold (p<0.05), with which a minimum score of 27 was used for peptide identification. The protein redundancy at the database under different accession numbers was limited to Mus musculus. All the proteins identified in the current study were found in these domains. The comparative study was performed using ProteoIQ v2.8 (local license) (Bonilla *et al*, 2023; Paez & Callegari, 2022).

### Gene Ontology (GO) enrichment analysis of the identified proteins

GO analysis of the proteins was carried out by using the DAVID functional annotation tool (https://david.ncifcrf.gov/summary.jsp) as previously described (Huber *et al*., 2022). For each protein, the UniProt ID was converted to its official gene ID, with Mus musculus selected as the species. Two genes per term and an ease threshold of 0.1 were utilized for the analysis (default settings). The p-value was converted to the -log(p). The differences in GO terms and pathways between control and *Psmc1* KO samples were determined using the Venny 2.1 software (https://bioinfogp.cnb.csic.es/tools/venny/).

### Western blot analysis

Western blot (WB) analysis was carried out as previously described methods (Liu *et al*, 2020; Liu *et al*, 2021). Briefly, the isolated cellular fractions or cortical lysates were diluted with a cell lysis buffer(Dong *et al*, 2012). An equal amount of protein was resolved on 12% SDS-PAGE gels, and the separated proteins were then transferred onto nitrocellulose membranes under standard conditions. The membrane was blocked in 5% nonfat milk in Tris-buffered saline supplemented with 0.5% Tween-20 and probed with the following primary antibodies: anti-Stat1 (1:1000, Cell Signaling), anti-Trem2 (1:1000, Cell Signaling), anti-Actin (1:1000, Developmental Studies Hybridoma Bank), anti-PSD95 (1:1000, Cell Signaling), anti-NF-κB (1:1000, Cell Signaling), and anti-GFAP (1:1000, Millipore). Following overnight incubation with the primary antibody, the blots were incubated with a secondary antibody for one hour. The secondary antibodies were conjugated with infrared dyes (1:3000, LI-COR, #926-32211; #92668072, #926-33214). Protein band intensities were quantified using ImageJ.

### Histopathological analysis

Brain fixation, sectioning, and immunohistochemical procedures were carried out using previously described protocols (Liu *et al*., 2021). Briefly, the brain sections were blocked in a blocking buffer (1% normal donkey serum, 0.1% Triton, and 0.05% Tween). Then, the samples were incubated with the following primary antibodies: anti-GFAP (1:1000, EMD Millipore) or anti-Iba1 (1:200, FUJIFILM Wako) overnight at 4°C. Next, the sections were incubated with a Cy3-conjugated goat anti-rabbit secondary antibody (1:200, Jackson ImmunoResearch). The nuclei were counterstained with the DNA-binding dye Hoechst 33342 (1:1000, Thermo Fisher). Images were collected using the Zeiss Axiovert Inverted Epifluorescent Microscope. Three sections per animal were analyzed. The analysis of GFAP and Iba1 consisted of counting the GFAP- or Iba1-positive cells before dividing them by the total (neuronal and glial) number of cells (Hoechst 33342-positive cells).

### Statistical analysis

GraphPad Prism Statistical Software version 9.4.1 was utilized for statistical analysis and graphical displays. All numerical data are presented as mean ± SEM. A student’s t-test was utilized to compare the two groups. For the analysis of the RAWM, average errors in finding the platform were calculated on a three-trial block. For comparisons between different groups of animals in the RAWM, repeated measures, and two-way ANOVA with Tukey’s post hoc test were utilized. Statistical significance was accepted when p < 0.05.

## Results

### Genetically disrupting the 26S proteasome reduces proteasome activity and impairs animal learning and memory

To understand the effect of 26S proteasome deficiency on brain function, we genetically ablated *Psmc1*, one 19S proteasomal subunit of the 26S proteasome (**Fig. 1A**), in the forebrain region. This caused the regulatory particle to be unable to bind to the 20S core particle, rendering the deficiency of the 26S proteasome in this brain region (Bedford *et al*., 2008; Ugun-Klusek *et al*, 2017). Utilizing this model, we initially examined the three proteasome activities, chymotrypsin-, caspase-, and trypsin-like proteasome activities, in the forebrain. As demonstrated in **Fig. 1B, 1C, and 1D**, all three of the proteasome activities were reduced in the KO brain tissues compared to the control animal tissues at two months of age. We then performed the RAWM test to assess their learning and memory capabilities. *Psmc1* KO mice exhibited a significant increase in the number of incorrect arm entries (errors) to find the platform compared to control animals at blocks 4, 5, 6, 7, 8, and 10, suggesting an impairment in learning and memory (**Fig. 1E**). A significant increase in the time taken by the mice to find the platform was also observed in the *Psmc1* KO animals compared to the control animals at blocks 1, 2, 4, 5, and 10 (**Fig. 1F**). It is noted that the differences between the control and *Psmc1* KO groups in block 10 appear greater than in blocks 5 to 9. This might be caused by the *Psmc1* KO mice being less tolerant to repeated swimming, causing them to make more errors at the late stage of the test. Thus, 26S proteasome deficiency impairs animal learning and memory.

### 26S proteasome deficiency causes synapse loss and neuroinflammation in the brain

Next, we assessed the synapses in the brain of mice at two months of age, as synapse loss exhibits the strongest correlation for cognitive decline in AD (de Wilde *et al*, 2016; DeKosky *et al*, 1996; Scheff & Price, 2006; Terry *et al*, 1991). We, therefore, isolated synaptosomes from the cortex of the *Psmc1* KO and the control animals to explore the possibility of synapse alterations. Synapse fractionation was confirmed by WB analysis of a postsynaptic and a presynaptic marker protein. As shown in **Fig. 2A and 2B**, the postsynaptic marker, PSD95, in the *Psmc1* KO synaptosomes, exhibited a significant reduction in expression compared to control synaptosomes, suggesting the possibility of post-synaptic loss occurring at this age; however, SNAP-25, a presynaptic marker, showed no differences between control and *Psmc1* KO animals in the synaptosome isolates at this stage (**Fig. 2A and 2C**).

**Fig. 2.**
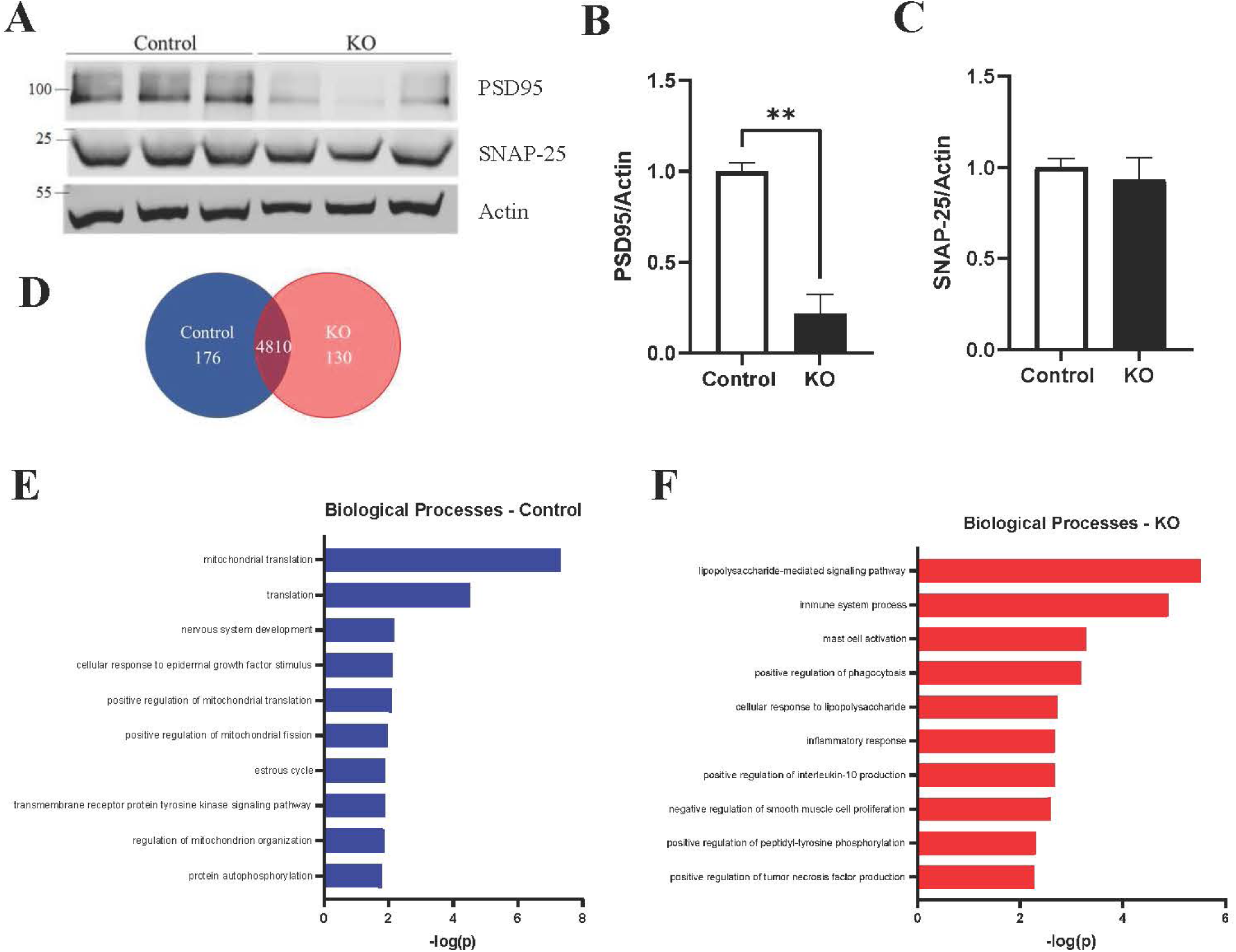
Mass spectrometry combined with bioinformatic analyses of synaptic proteins identifies inflammation and immune responses as the top listed pathways in *Psmc1* KO brains at two months of age. Synaptosomes were isolated from the cortex of control and *Psmc1* KO animals. *Psmc1* KO animals showed a significant decrease in PSD95, a postsynaptic marker, compared to control animals (**A and B**). SNAP-25, a presynaptic marker, exhibited no statistical difference between control and *Psmc1* KO animals (**A and C**). Data are presented as mean ± SEM; n = 3 per genotype, ** p < 0.01 (Data was analyzed by a student’s t-test). Actin was used as a loading control. Mass spectrometric analysis of synaptic proteins revealed 176 proteins unique to control synaptosomes and 130 proteins unique to *Psmc1* KO synaptosomes (**D**). GO enrichment analysis for the top biological processes unique to control (**E**) and *Psmc1* KO synaptosomes (**F**) (n = 3 per genotype, two females and one male).

With the isolated synaptosomes, we then performed proteomic analysis using the mass spectrometry (MS)-based approach, which is an unbiased method commonly used for proteomics. MS analysis of synaptosomal proteins in the control samples revealed 176 unique, diverse proteins. Surprisingly, only 130 proteins were found exclusively in *Psmc1* KO synapses (**Fig. 2D**). Gene Ontology (GO) analysis of control and *Psmc1* KO synaptosomal proteins identified differential molecular functions between them. The biological processes from control synaptosomes identified processes associated with mitochondrial function and nervous system development (**Fig. 2E**). However, the *Psmc1* KO synaptosomes exhibited biological processes associated with the immune system response and inflammatory response (**Fig. 2F**), providing further support for the influence of proteasome dysfunction on neuroinflammation. The molecular functions identified in control synaptosomes consisted of the structural constituent of the ribosome and RNA binding as the top two pathways (data not shown). In contrast, *Psmc1* KO synaptosomes consisted of molecular functions associated with inflammation and transmembrane signaling receptor activity (data not shown), providing further support for an inflammatory response occurring at the synapse. This data suggests that *Psmc1* KO synaptosomes contain proteins associated with inflammation that are absent from control synaptosomes.

### 26S proteasome deficiency causes the accumulation of inflammation-associated proteins

As inflammation is a biological response that is not limited to synapses, we next examined some of the identified inflammation-related proteins, including Stat1, Trem2, and NF-κB in the cortex of control and *Psmc1* KO brain lysates using WB analysis. Stat1and NF-κB are transcription factors and activation of these two pathways are the most common inflammatory signaling pathways following immune responses (Chen *et al*, 2018). Trem2 is an innate immune cell surface receptor regulating microglial function and is involved in the pathogenesis of many neurodegenerative diseases (Diaz-Lucena *et al*, 2021; Yang *et al*, 2020). Recent genome-wide association studies identified rare variants in the gene-triggering receptor expressed on myeloid cells 2 (*Trem2*) that are associated with a high risk of developing late-onset AD (Guerreiro *et al*, 2013; Jonsson *et al*, 2013; Neumann & Daly, 2013). In our study, Stat1 expression significantly increased in *Psmc1* KO brain samples compared to control samples (**Fig. 3A and 3B**). Similarly, we observed a significant increase in Trem2 and NF-κB in *Psmc1* KO brain samples compared to control samples (**Fig. 3A-3D**). It is noted that two bands were seen in NF-κB blotting; the lower band represents the mature form, while the top band represents the precursor. NF-κB consists of five related proteins and can exist in the form of dimers, either homo- or heterodimers, to bind to DNA. P65 forms a dimer with p50. In the canonical pathway, p65/p50 dimers are in an inactive state sequestered in the cytoplasm; however, when exposed to proinflammatory stimuli, IκBα releases the p65/p50 dimers, where they can translocate to the nucleus to bind to the cognate κB motif to activate NF-κB target genes (Collins *et al*, 2016).

**Fig. 3.**
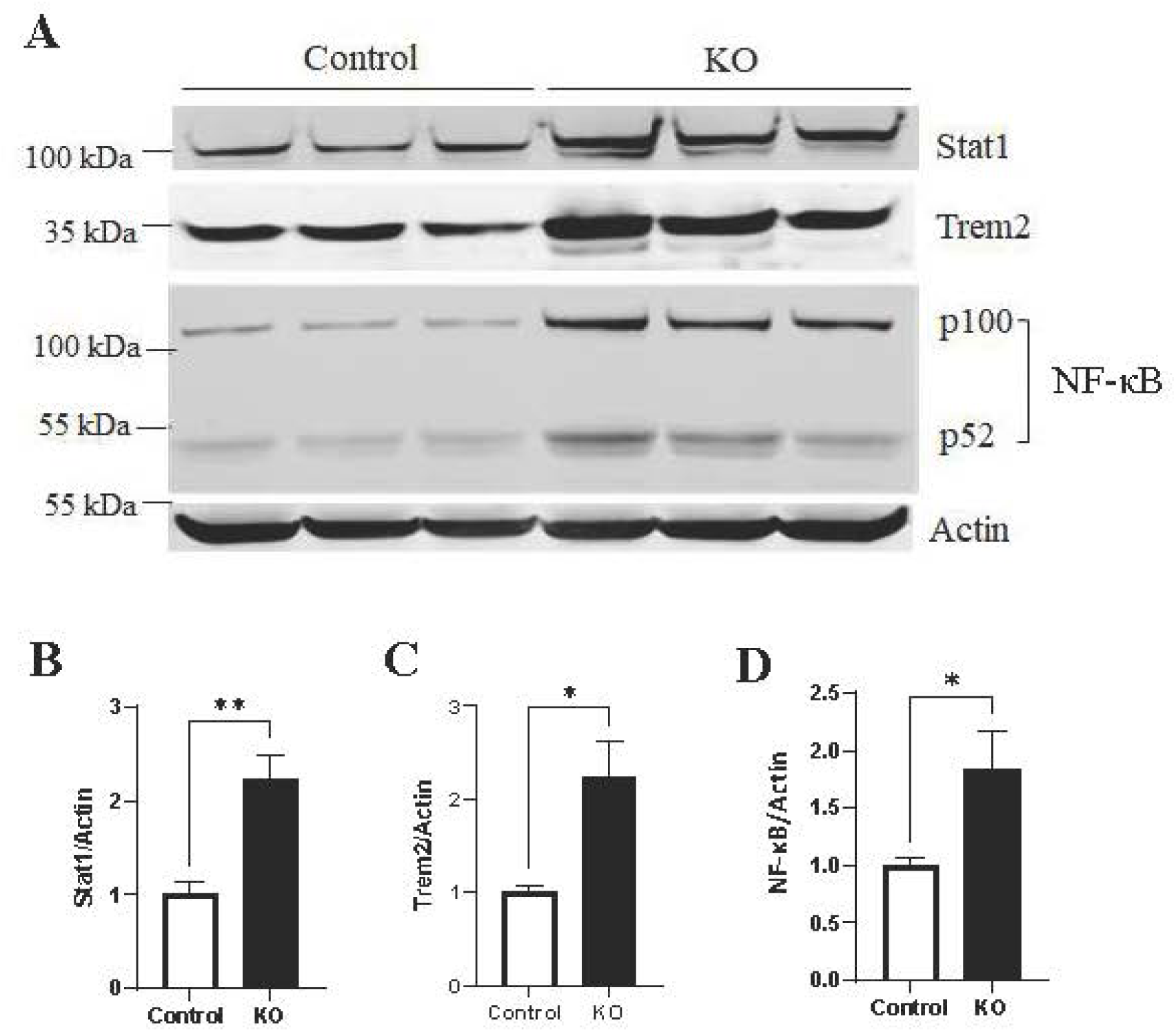
26S proteasome deficiency causes neuroinflammation at two months of age. WB analysis was conducted to define the expression of Stat1, Trem2 and NF-κB in *Psmc1* KO and control brain samples (**A**). Quantitation of inflammatory-associated protein levels for Stat1 (**B**), Trem2 (**C**), and NF-κB (**D**). Actin was used as a loading control. Data are presented as mean ± SEM; n = 6 per genotype, * p < 0.05; ** p < 0.01 (Data analyzed by a student’s t-test).

Therefore, 26S proteasome deficiency upregulates key inflammatory regulatory proteins, including NF-κB, a master regulator of the innate immune response (Dorrington & Fraser, 2019).

### 26S proteasome deficiency activates astrocytes and microglia

As activation of microglia and astrocytes is a common feature of neuroinflammation and is involved in several neurodegenerative diseases (Kwon & Koh, 2020), we next assessed whether the two types of glia were activated. The glial fibrillary acidic protein (GFAP), a protein indicative of astrocytes, represents a biomarker of neuroinflammation, as an increase in its expression of GFAP indicates more of a reactive phenotype of astrocytes (Wilhelmsson *et al*, 2006). WB analysis of GFAP protein levels in the brains of controls and *Psmc1* KO samples indicated a significant increase in GFAP in *Psmc1* KO samples compared to controls, providing further support for overt neuroinflammation occurring in the brain following 26S proteasome disruption (**Fig. 4A and 4B**).

**Fig. 4.**
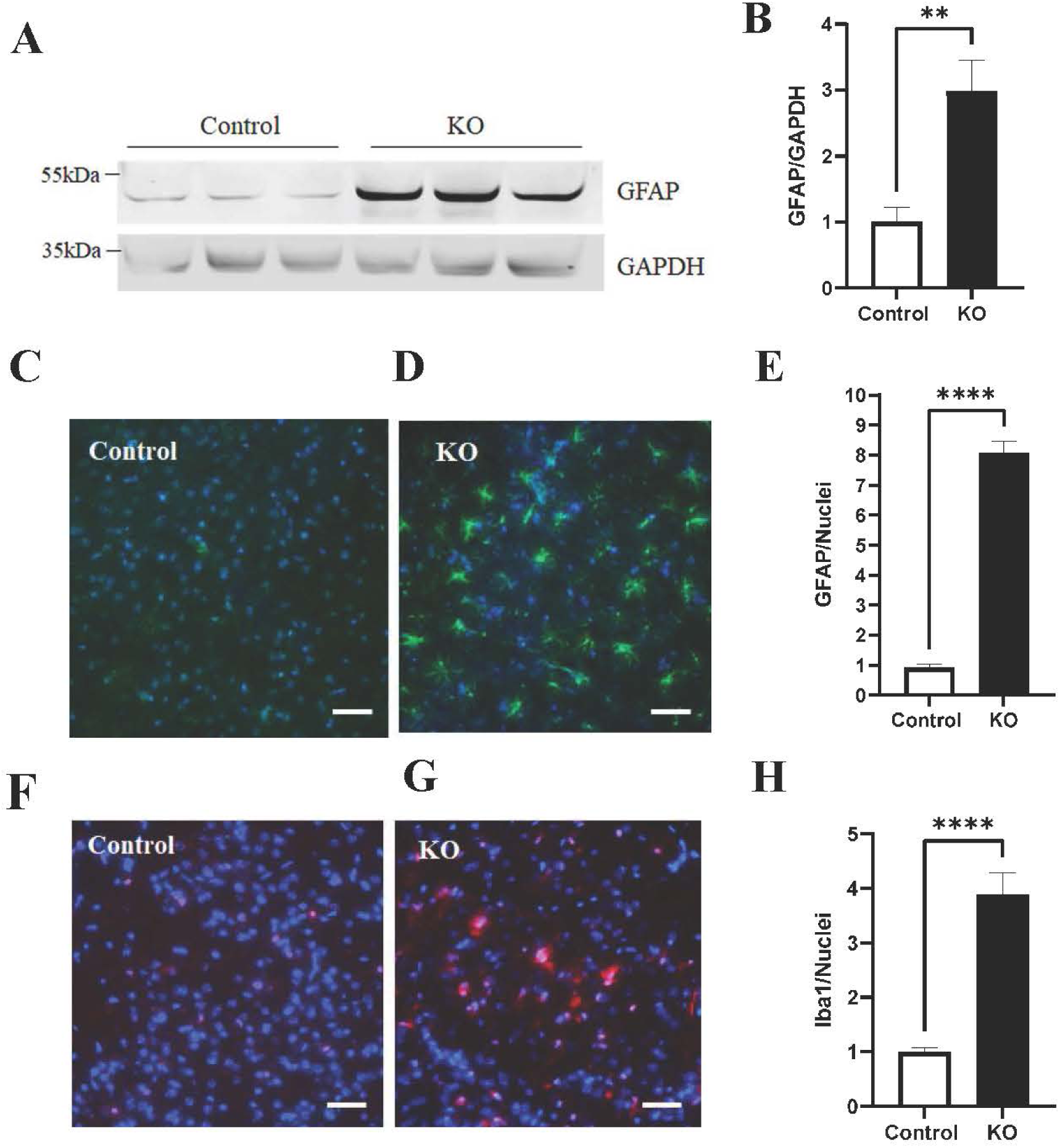
26S proteasome deficiency activates astrocytes and microglia at two months of age. WB analysis of GFAP (**A and B**) in the cortex of *Psmc1 KO* and control samples. Immunohistochemical analysis of the cortex of GFAP (**C to E**) and Iba1 (**F to H**) suggests proteasome disruption causes an overt inflammatory response. Scale bar, 50 µm. Data are presented as mean ± SEM; n = 3 – 6 per genotype, one to three females and two to three males for each genotype; * p < 0.05; ** p < 0.01, *** p < 0.001, **** p < 0.0001 (Data analyzed by a student’s t-test; blue, Hoechst; green, GFAP; red, Iba1).

Immunohistochemistry analysis of GFAP revealed significantly more GFAP staining in the cortex of *Psmc1* KO brains compared to control samples, suggesting reactive gliosis occurring following 26S proteasome disruption (**Fig. 4C-4E**). Ionized calcium-binding adaptor molecule 1 (Iba1) is a marker for microglia, the resident macrophage cell in the brain (Guan *et al*, 2022). An increase in Iba1 expression is indicative of microglial activation (Hovens *et al*, 2014). There was a significant increase in Iba1-positive cells in the brains of *Psmc1* KO animals compared to control samples, providing further support for an inflammatory response occurring in the brains of *Psmc1* KO animals (**Fig. 4F-4H**).

### Inhibition of NF-κB improves learning and memory in *Psmc1* KO animals

The aforementioned results consistently support that disrupting the 26S proteasome causes neuroinflammation and activation of the NF-κB pathway. As NF-κB is a master regulator of innate immune responses and plays a crucial role in modulating inflammatory responses (Dorrington & Fraser, 2019; Sun *et al*, 2022), we assessed whether treatment of Psmc1 KO mice with PDTC, a selective and blood-brain barrier-permeable NF-κB inhibitor (Kan *et al*, 2016), confers any beneficial effect on the learning and memory of mice. We, therefore, treated the KO mice at five weeks of age and then continued the treatment for three weeks with daily intraperitoneal injections of 50 mg/kg of PDTC (Qin *et al*, 2014) prior to behavioral tests at seven weeks. Mice were sacrificed at eight weeks for examination of inflammation (**Fig. 5A**). Behavioral tests for PDTC-treated animals began when the mice reached seven weeks of age (**Fig. 5A**). The learning and memory capability of animals was assessed using the RAWM following NF-κB inhibition. *Psmc1* KO + PDTC-treated animals demonstrated better learning and memory capability compared to the KO control animals at every block. At blocks 3, 4, 8, 9, and 10, the *Psmc1* KO + PDTC-treated animals performed significantly better than the KO control animals (p < 0.05). The *Psmc1* KO + PDTC-treated animals performed similarly to the non-KO control mice (Control) on blocks 8-10 (p > 0.05) (**Fig. 5B**). Furthermore, *Psmc1* KO + PDTC-treated animals spent less time finding the platform in the RAWM compared to the KO control mice (p < 0.05) (**Fig. 5C**). This data suggests that NF-κB inhibition can reverse the learning and memory impairments caused by 26S proteasome disruption.

**Fig. 5.**
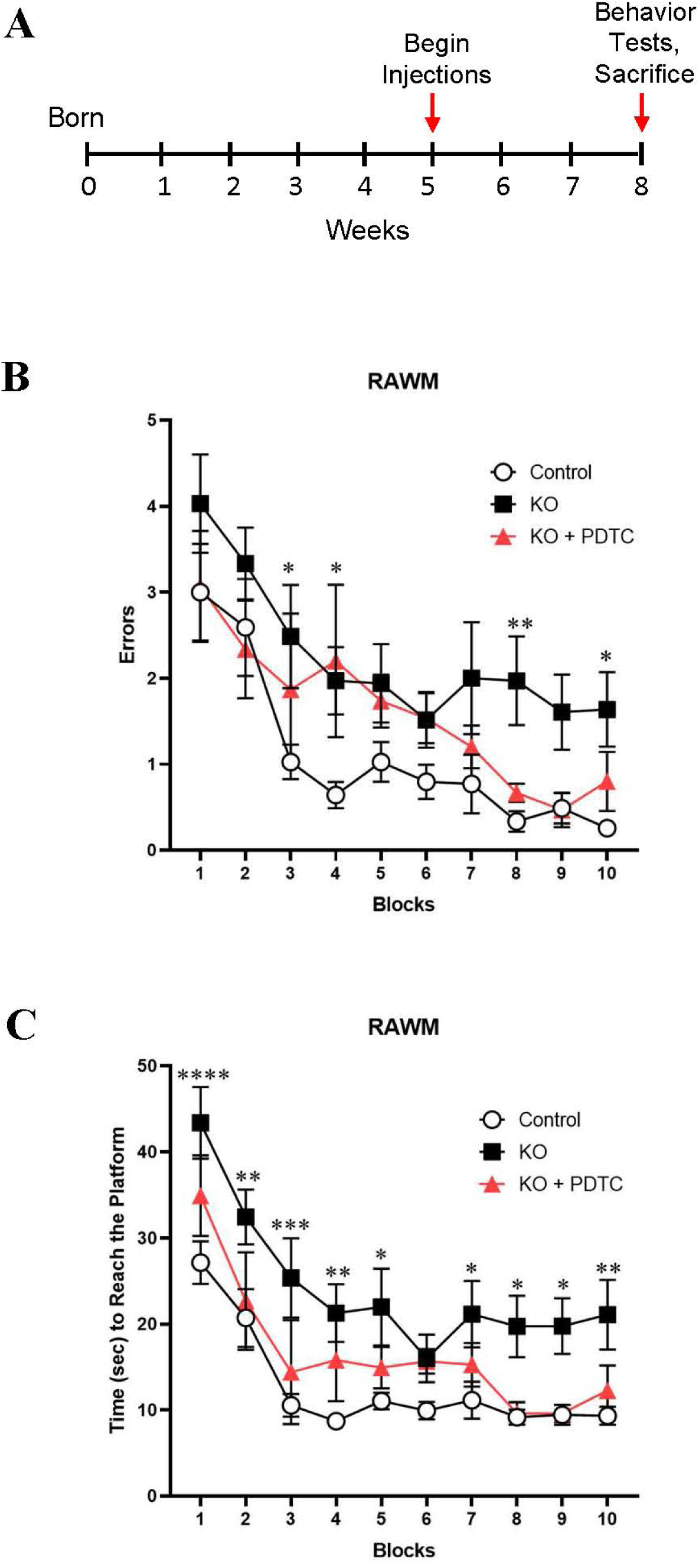
Inhibition of NF-κB improves learning and memory in *Psmc1* KO mice. Mice were treated with the NF-κB inhibitor, PDTC, at five weeks of age for three weeks (**A**). Diagram illustrating experimental paradigm (**A**). The RAWM was used to assess animals’ learning and memory capabilities (**B**) and (**C**). Data are presented as mean ± SEM; n = 5 – 29 animals per genotype, 10 females and 19 males for the “Control” group, six females and five males for “KO” group, two males and three females for “KO + PDTC” group; * p < 0.05; ** p < 0.01. RAWM data were analyzed by two-way ANOVA and Tukey’s post hoc test.

### Inhibition of NF-κB improves 26S proteasome deficiency-caused neuroinflammation

As we observed increased inflammation-associated proteins in the brains of *Psmc1* KO mice, we next examined the effect of NF-κB inhibition on the expression of Stat1 and Trem2. To this end, we performed a WB analysis of the proteins following the drug treatment. Stat1 expression significantly decreased when *Psmc1* KO animals were treated with the NF-κB inhibitor compared to the KO control samples (**Fig. 6A and 6B**). We found a significant decrease in Trem2 in *Psmc1* KO + PDTC treated samples compared to the KO control samples (**Fig. 6A and 6C**). These results indicate that NF-κB inhibition suppresses 26S proteasome deficiency caused upregulation of Stat1 and Trem2 in *Psmc1* KO animals.

**Fig. 6.**
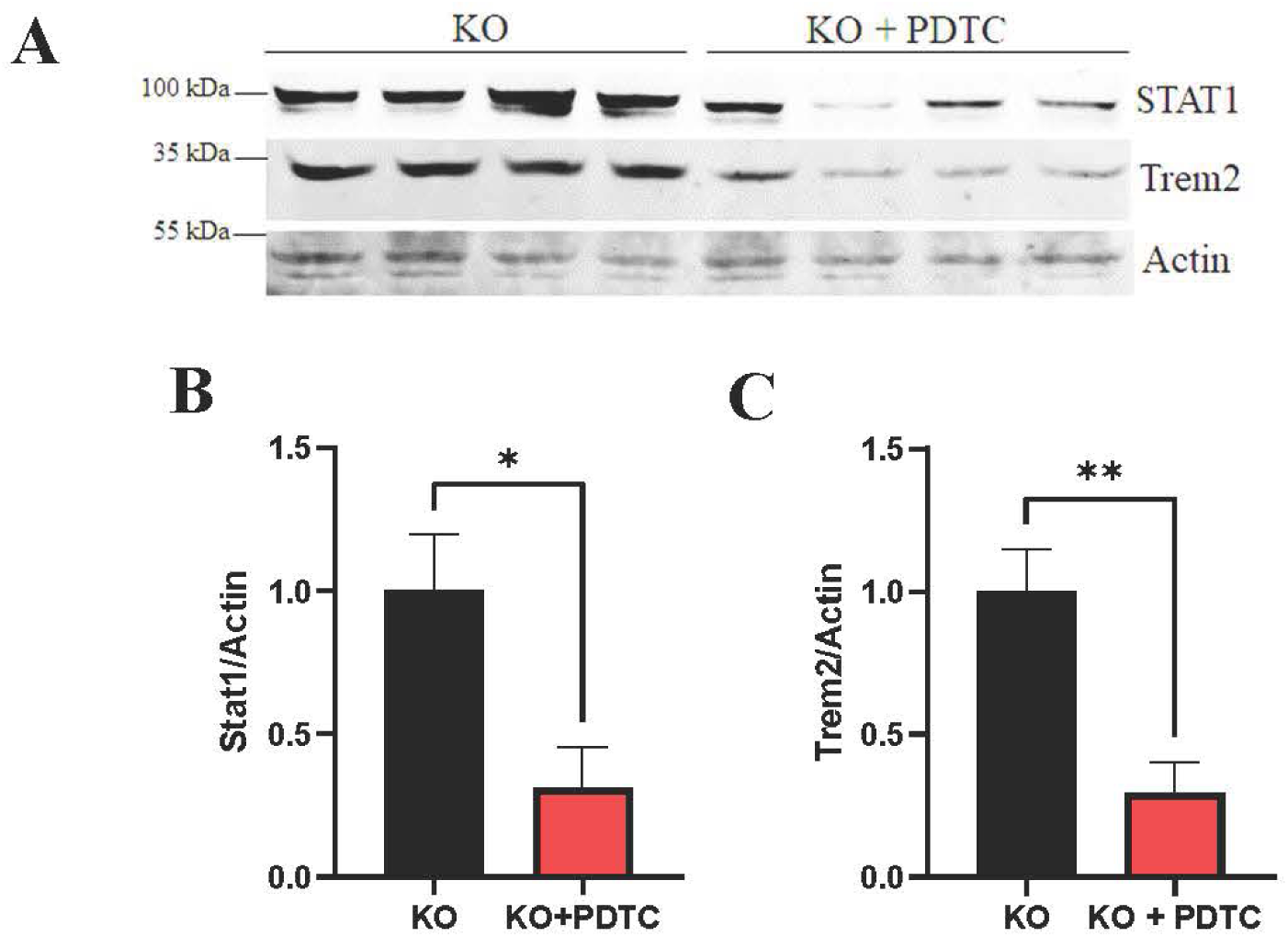
Inhibition of NF-κB attenuates neuroinflammation in *Psmc1* KO mice. WB analysis of the expression of Stat1 and Trem2 in *Psmc1* KO and control brain samples revealed that NF-κB inhibition decreased the level of Stat1 (**B and C**) and Trem2 (**B and D**) in the brains of *Psmc1* KO animals. (**A**). The representative ratio of inflammatory-associated proteins compared to Actin (loading control) for Stat1 (**C**), and Trem2 (**D**). Data are presented as mean ± SEM; n = 3 – 4 per genotype, * p < 0.05; ** p < 0.01 (Data analyzed by a student’s t-test).

### Inhibition of NF-κB suppresses activation of astrocytes and microglia in the KO mouse brains

Immunohistochemistry analysis of GFAP revealed significantly reduced GFAP in the cortex of *Psmc1* KO + PDTC treated mice compared to the KO control animals, suggesting blockage of reactive astrocytes following the inhibition of NF-κB (**Fig. 7A-7C**). Similarly, there was a significant decrease in Iba1-positive cells in the brains of *Psmc1* KO + PDTC treated mice compared to the KO control animals (**Fig. 7D-7F**), providing further support for the attenuation of the inflammatory response in *Psmc1* KO animals following NF-κB inhibition.

**Fig. 7.**
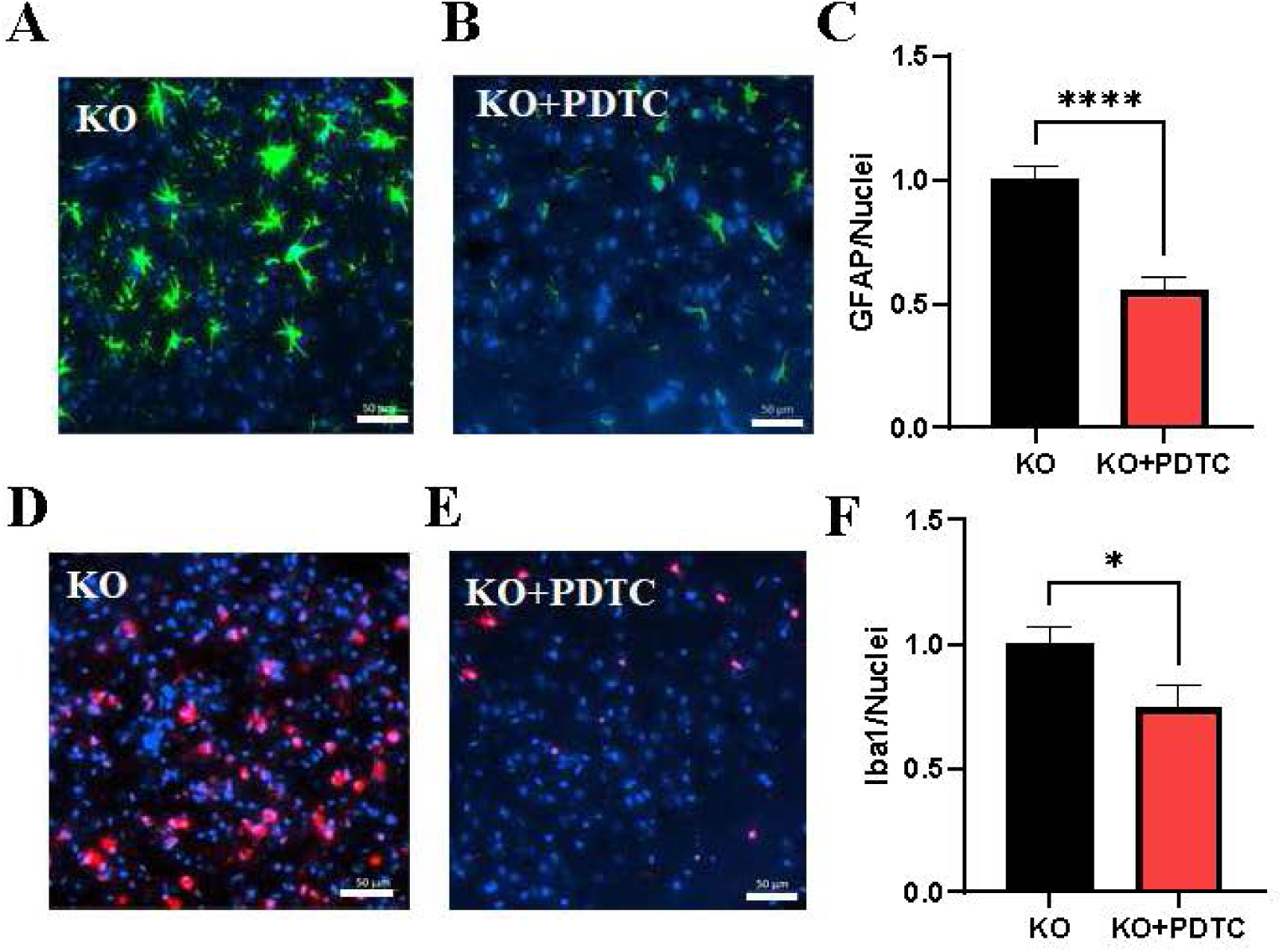
Inhibition of NF-κB suppresses activation of astrocytes and microglia in *Psmc1* KO mouse brain. Immunohistochemical analysis of the cortex of GFAP (**A-C**) and Iba1 positively stained cells (**D-F**). Scale bar, 50 µm. Data are presented as mean ± SEM; n = 3 per genotype (two males and one female for each genotype), * p < 0.05 and **** p < 0.0001 (Data analyzed by a student’s t-test; blue, Hoechst; green, GFAP; red, Iba1).

## Discussion

Impaired proteasome and neuroinflammation have been implicated in various neurodegenerative disorders, including AD. However, whether impaired proteasome is a causative mechanism to neuroinflammation is less understood. Here, we took advantage of a genetic mouse model to ablate the function of the 26S proteasome that is primarily responsible for ubiquitin-dependent protein degradation to determine the effect of disrupting 26S proteasome function on neuroinflammation and neuronal function. Our results consistently indicated that 26S proteasome deficiency caused synapse loss, neuroinflammation, and learning and memory dysfunction. Importantly, we identified that 26S proteasome deficiency-caused neuroinflammation was mediated largely by the NF-κB signaling pathway. Hence, our data provide direct evidence that impaired 26S proteasome causes neuroinflammation in the brain.

In addition to neuroinflammation, loss *of* synapses is another key pathological hallmark of AD. Our proteomics data revealed a reduction of total number of synaptic proteins, which may be caused by loss of synapses or decreased production of synapses due to neuroinflammation. Intriguingly, in an AD transgenic mouse model, it was found that there was more loss of postsynaptic proteins than presynaptic proteins (Gylys *et al*, 2004). In accordance with this finding, we observed a diminished level of PSD95, a postsynaptic marker, from the isolated synaptosomes from the *Psmc1* KO mouse brains, indicating the possible loss of synapses. Previous data indicates that reduced PSD95 protein levels can alter neuronal functions, consequently leading to synaptic loss and neurodegeneration (Sultana *et al*, 2010). Interestingly, SNAP25, a presynaptic marker, did not decrease in the isolated synaptosomes from the *Psmc1* KO brain at this stage. Further work is necessary to determine if synaptic loss occurs at later time points. Neuroinflammation observed in the *Psmc1* KO mouse brain may account for the reduced PSD95 because some inflammation-related transcription factors, such as Stat1, are responsible for impairing the structure and function of synapses (Han *et al*, 2023; Hsu *et al*, 2014; Li *et al*, 2019b; Ma *et al*, 2021).

Our data consistently support the significant role of NF-κB, a master regulator of innate immune responses (Dorrington & Fraser, 2019; Gu *et al*, 2015; Khalil *et al*, 2019; Sun *et al*., 2022) in 26S proteasome deficiency-induced neuroinflammation. First, our proteomics and WB results showed that NF-κB levels were increased in *Psmc1* KO mouse brains. Second, our transcriptome analysis identified the NF-κB signaling pathway as one of the top listed pathways among the upregulated genes following proteasome disruption. Third, inhibition of NF-κB was able to attenuate 26S proteasome deficiency-caused neuroinflammation in *Psmc1* KO mouse brain and reduce learning and memory impairment. These data suggest that 26S proteasome deficiency-caused neuroinflammation is mediated, at least partially, through the NF-κB signaling pathway (**Fig. 8**). Activated NF-kB then induces neuroinflammation via Stat1, Trem2, and other downstream factors, leading to synapse alterations and impaired cognitive function (**Fig. 8**).

**Fig. 8.**
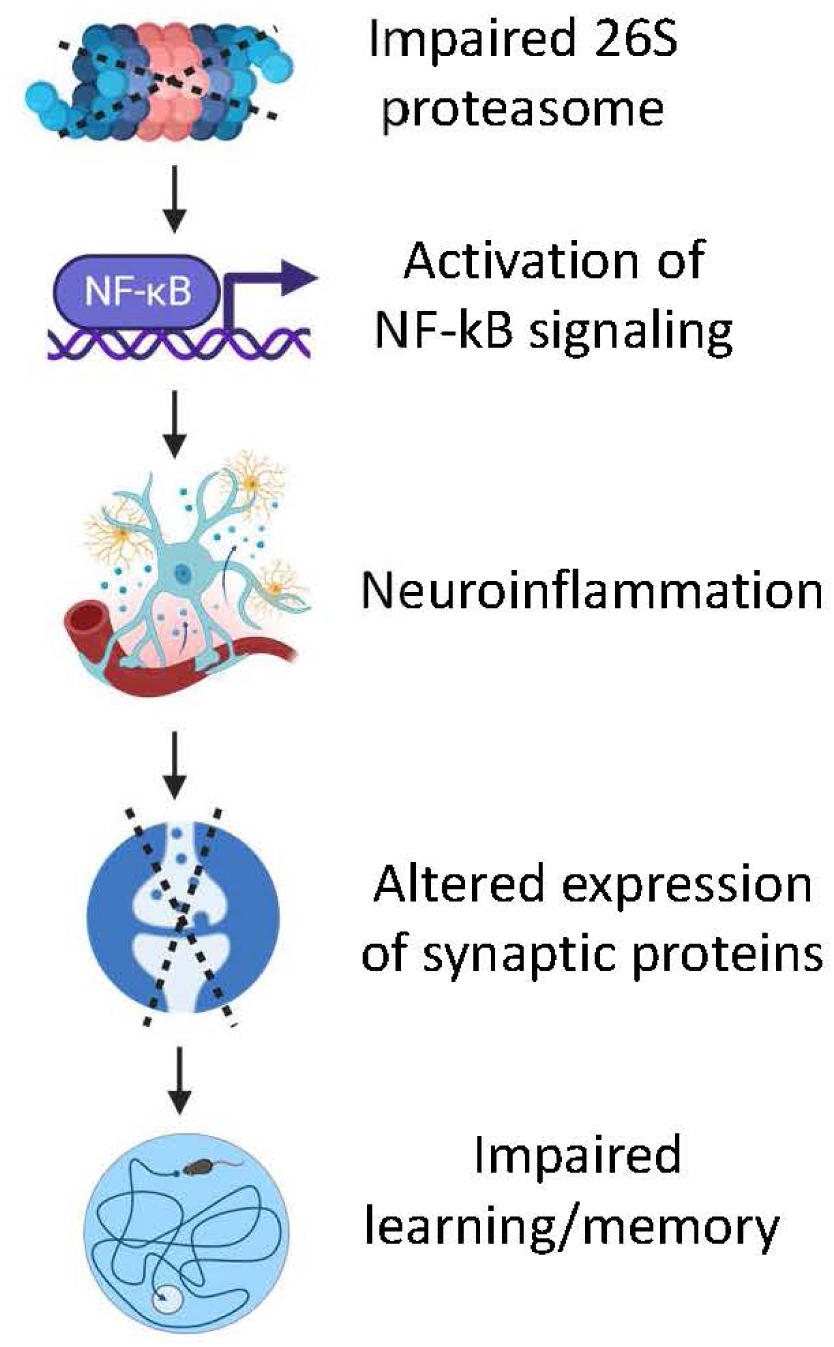
Working model. Impaired 26S proteasome causes neuroinflammation, synaptic alterations, and disrupted learning and memory via activation of NF-kB.

As NF-κB can be degraded by the 26S proteasome (Liu & Chen, 2011), impaired proteasome would result in increased NF-κB. As a master regulator of immune responses, NF-κB induces the expression of diverse proinflammatory genes, including those encoding cytokines and chemokines (Sun *et al*., 2022). A high NF-κB level is correlated with a high level of β-site amyloid precursor protein (APP) cleaving enzyme-1 (BACE), and the p65 subunit of NF-κB can bind to the κB elements on the promoter of BACE to facilitate the expression of β-secretase (Chen *et al*, 2012). This results in the cleavage of APP into Aβ, leading to the formation of Aβ fibrils. Additionally, Aβ oligomers can activate NF-κB (Chen *et al*., 2012), astrocytes, and microglia (Di Benedetto *et al*, 2022). Therefore, 26S proteasome disruption may induce a chain of reactions, involving multiple mechanisms, which cause NF-κB levels to become even higher, subsequently leading to increased production of proinflammatory cytokines to exacerbate activation of astrocytes and microglia that causes elevated levels of GFAP and Iba1. Moreover, we found that other immune response-related proteins, Stat1 and Trem2, also increased in the *Psmc1* KO mouse brain. NF-κB regulates Trem2 to shape innate immune and phagocytic responses by microglia to contribute to neuroinflammation (Zhao *et al*, 2013). The increase of Trem2 observed in this study might be due to a negative feedback or compensation effect of inflammation to curb NF-κB-induced neuroinflammation because prior data have shown that Trem2 inhibits neuroinflammation via downregulating NF-κB signaling in microglia (Li *et al*, 2019a).

Although Stat1 is also upregulated in the KO mouse brain, it may not be a targeted gene of NF-κB, as the two transcriptional factors are activated by distinct extracellular stimuli and may need to cross-talk to confer a synergy in regulating the expression of inflammatory genes at the transcriptional level (Barbosa Lima *et al*, 2019). Conversely, inhibition of NF-κB reduced Stat1 level should be neuroprotective in the Psmc1 KO mouse brain (Han *et al*., 2023). These distinct mechanisms work together, contributing to neuroinflammation observed in mice.

In humans, the variance of *Psmc1* is associated with an autosomal recessive syndrome of severe developmental delay and intellectual disability, hearing loss, and other problems (Aharoni *et al*, 2022). Impaired learning and memory observed from *Psmc1* KO mice are consistent with this clinical data, indicating that impaired proteasome could be a driver for the development of an AD-like phenotype. This study demonstrates that proteasome function and neuroinflammation may go “hand-in-hand,” as neuroinflammation was exacerbated in 26S proteasome deficiency mice. Indeed, this possibility is strengthened by our data that inhibiting one inflammatory signaling pathway, NF-κB, can reverse the phenotype-induced by 26S proteasome disruption.

## Conclusion

In this study, we discovered that 26S proteasome deficiency impairs learning and memory and causes neuroinflammation. We also identified a significant mediator, NF-κB, in modulating the 26S impairment-caused neuroinflammation and neurodysfunction. Our results will help better understand the pathogenesis of AD and other neurodegenerative disorders where impaired proteasome is consistently coupled with neuroinflammation.

## Ethics statement

All procedures used to handle mice were approved by the Institutional Animal Care and Use Committee at the University of South Dakota and Texas Tech University Health Science Center, and were in accordance with the National Institute of Health Guide for the Care and Use of Laboratory Animals.

## Author Contributions

HW conceived the presented idea and model. HW, CCH, and EC designed the experiments, and CCH, EC, MP, and XL performed experiments and analyzed the results. CCH and HW interpreted the data and wrote the manuscript. AD edited the manuscript.

## Acknowledgments

We would like to thank Drs. R. John Mayer and Zubair Karim for providing the floxed *Psmc1* mouse model. We also express our gratitude to the SD-BRIN Proteomics Core at the University of South Dakota for the technical assistance in proteomics and also to Ryan Johnson and Bill Conn from the University of South Dakota Information Technology Research Computing for their help in server operation and maintenance.

